# Relationship between salmon egg subsidy and the distribution of an avian predator

**DOI:** 10.1101/2022.07.19.500590

**Authors:** Taihei Yamada, Hirotaka Katahira, Kazuki Miura, Futoshi Nakamura

## Abstract

As a spatial subsidy, which is the phenomenon of transferring resources from a donor system to a recipient system, anadromous salmonids contribute to the supply of marine-derived nutrients to freshwater and terrestrial systems. Live salmon and salmon carcasses and eggs are utilized by various organisms and affect their abundance and distribution. However, the evaluation of the effect of salmon subsidies on the abundance and distribution of terrestrial animals is biased towards predators or scavengers that utilize spawning adults and carcasses, and few studies have focused on the effect of salmon eggs as a subsidy. To avoid underestimating the function of salmon subsidies, the response to the availability of salmon eggs in various systems should be investigated. Here, we investigated the abundance and feeding behaviour of the brown dipper *Cinclus pallasii*, as salmon egg a consumer, based on the hypothesis that the availability of salmon eggs affects the diet composition and stream distribution of this small predator. In addition, to test whether changes in the abundance of brown dippers are determined by salmon spawning, their abundance was compared upstream and downstream of the dam. Brown dippers used salmon eggs during the spawning season (53.7% of diet composition), and their abundance increased as the number of spawning redds increased. In contrast, this pattern was not observed upstream of the dam. These results suggested that the abundance and stream distribution of brown dippers vary according to the variation in the spatiotemporal availability of salmon eggs.

## 1. INTRODUCTION

Spatial subsidies are a phenomenon in which resources are transferred from a donor system to a recipient system (Polis et al., 1997). Spatial subsidies play a crucial role in biological communities because they affect the abundance and distribution patterns of organisms and the food web structure in the recipient systems by affecting the availability of basal resources (Hocking et al., 2013; Kawaguchi et al., 2003; Nakano & Murakami, 2001; Spiller et al., 2010; Terui et al., 2018).

Anadromous salmonids, well-known spatial subsidy representatives, transport marine-derived nutrients and energy to freshwater and terrestrial ecosystems through their migrations (Gende et al., 2002; Hocking & Reynolds, 2011; Koshino et al., 2013; Schindler et al., 2003). Salmon subsidies contribute to increasing aquatic invertebrate biomass and freshwater fish abundance in rivers (Denton et al., 2009; Wipfli et al., 1998, 1999). Spawning adults and carcasses are also used as food by terrestrial animals not only in the underwater ecosystem but also in surrounding riparian ecosystems and eventually affect the abundance and distribution of terrestrial scavengers and top predators (Boulanger et al., 2004; Christie & Reimchen, 2005; Field & Reynolds, 2013; Levi et al., 2012; Walters et al., 2021). Salmon subsidies thus provide insights into how multiple ecosystems are tangled together.

Past investigations on the effects of salmon subsidies on terrestrial organism abundance and distribution have thus far been biased towards predators or scavengers that utilize spawning adults and carcasses (e.g., Boulanger et al., 2004; Christie & Reimchen, 2005; Field & Reynolds, 2013; Levi et al., 2012; Walters et al., 2021). Considering that terrestrial organisms are provided multiple resources by salmon runs, such as eggs and fry, as well as spawning adults and carcasses (Munro, 1941; Willson & Halupka, 1995), salmon subsidies may have even more unexpected far-reaching effects. Therefore, a comprehensive understanding of the effect of subsidies on the population and distribution of organisms is important. However, few studies have focused on the effect of salmon eggs as a subsidy on the abundance and distribution of terrestrial animals. It is necessary to clarify the response to the availability of salmon eggs in various systems to avoid underestimating the function of salmon subsidies.

Dippers (Aves: Cinclidae) are riparian birds that mainly feed on aquatic invertebrates by diving into the water (Eguchi, 1990; Taylor & O’Halloran, 1997, 2001) and which are known to use salmon eggs in available rivers and seasons (Goodge, 1959; Obermeyer et al., 1999, 2006; Reimchen, 2017; Whitehorne, 2010). For example, the American dipper *Cinclus mexicanus* can achieve higher reproductive success (as measured by fecundity and juvenile growth) in reaches where *Oncorhynchus* swim upstream than in reaches where it does not (Obermeyer et al., 2006; Tonra et al., 2016). The population size of the white-throated dipper *C. cinclus* in Norway may also benefit from eating salmon fry because it was correlated with the annual density of salmon fry (Nilsson et al., 2018). Because dippers, which are not scavengers, are not affected by the amount of carcasses – in addition to their well-studied relationship with salmon, as noted above – they are a suitable model species for examining the effect of salmon egg subsidies on the abundance and distribution of terrestrial animals. The brown dipper *C. pallasii* (Figure 1), which is distributed in Asia (Hong et al., 2019), preys on salmon eggs and juvenile salmon (Murata, 1900). However, its actual status has never been evaluated quantitatively, and the relationship with salmon subsidies has long been overlooked.

**Figure 1.**
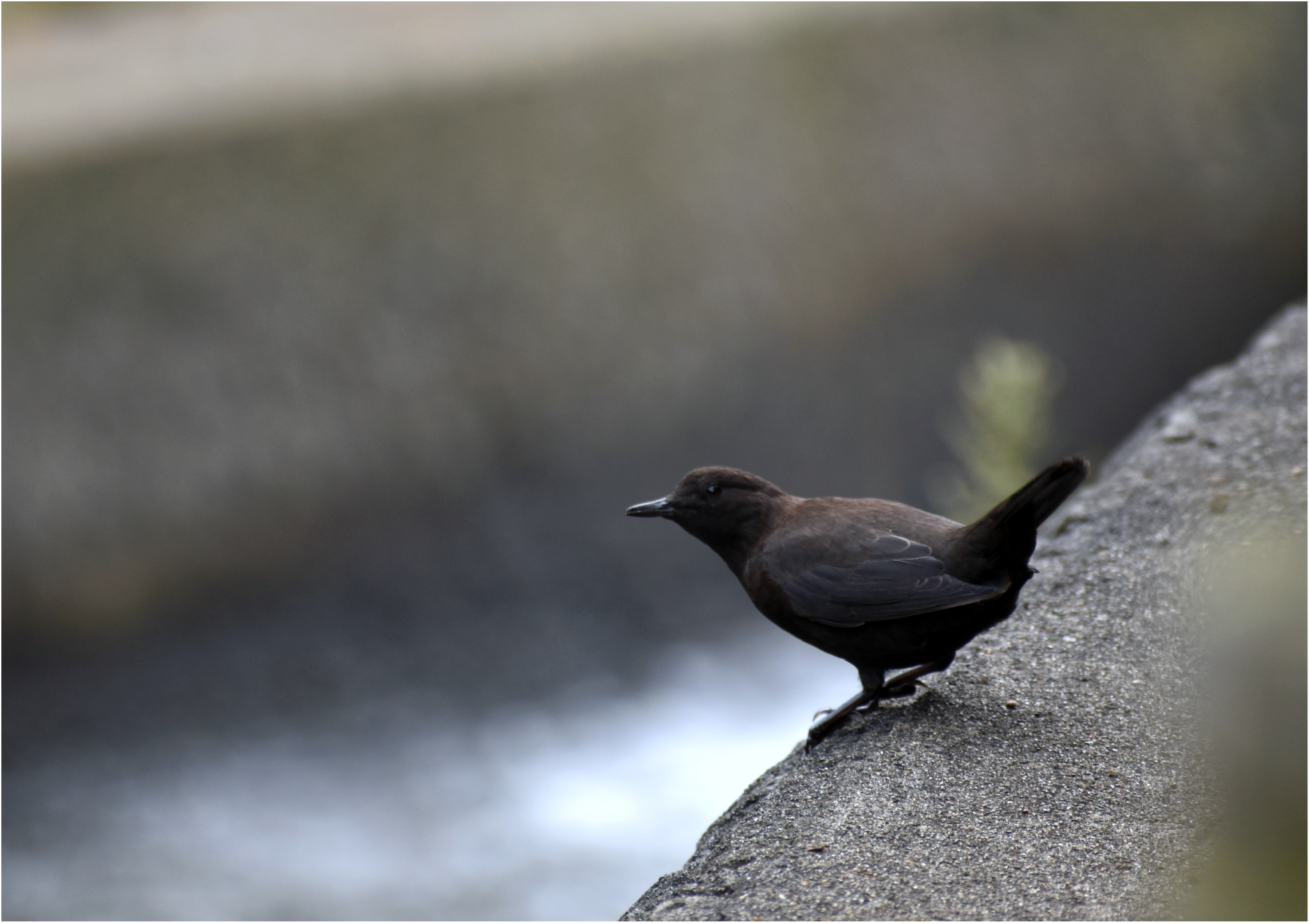
The brown dipper *Cinclus pallasii*. Photo by Yuya Eguchi.

We therefore investigated the abundance and diet composition of the brown dipper in the Shiretoko Peninsula of Hokkaido, northern Japan, where the spawning migrations of salmon are well observed, based on the hypothesis that the availability of salmon subsidies drives the diet composition and stream distribution of this small predator. In addition, by comparing the abundance of brown dippers above and below dams where salmon cannot run upstream during peak salmon spawning runs, we tested whether changes in the abundance of brown dippers are determined by salmon spawning. More specifically, it was predicted that salmon spawning would cause a shift in the diet of brown dippers to salmon eggs and an increase in the abundance of brown dippers by altering the distribution of food resources, while no such pattern occurs upstream of the dam.

## 2. MATERIALS AND METHODS

### 2-1. Study site

Four streams located in the Shiretoko Peninsula were selected for the present survey (Figure 2). Natural spawning sustains pink salmon *Oncorhynchus gorbuscha* and chum salmon *O. keta* populations in these streams, and the former is dominant (T. Yamada, unpublished data). The release of juvenile chum salmon has been conducted only in the Mosekarubetsu stream, and in-stream harvesting does not occur in all streams. The central part of the Shiretoko Peninsula has been designated as a World Natural Heritage site since 2005, partially because of the close relationships between the marine and terrestrial ecosystems sustained by the anadromous migration of pink salmon and chum salmon (IUCN, 2005). The upper reaches of the studied streams are included in the Shiretoko World Natural Heritage site. Rivers and streams in the Shiretoko Peninsula are highly fragmented by more than 330 artificial dams (Takahashi et al., 2005), which is no exception in all selected streams. Study sections were set up in each stream from the mouth of the stream to the proximal dam. The Chienbetsu stream was surveyed up to the second proximal dam because many salmon pass through the first proximal dam. The length of the study section was 289.4, 348.2, 154.9, and 211.3 m in the Chienbetsu stream, Funbe stream, Mosekarubetsu stream, and Shoji stream, respectively.

**Figure 2.**
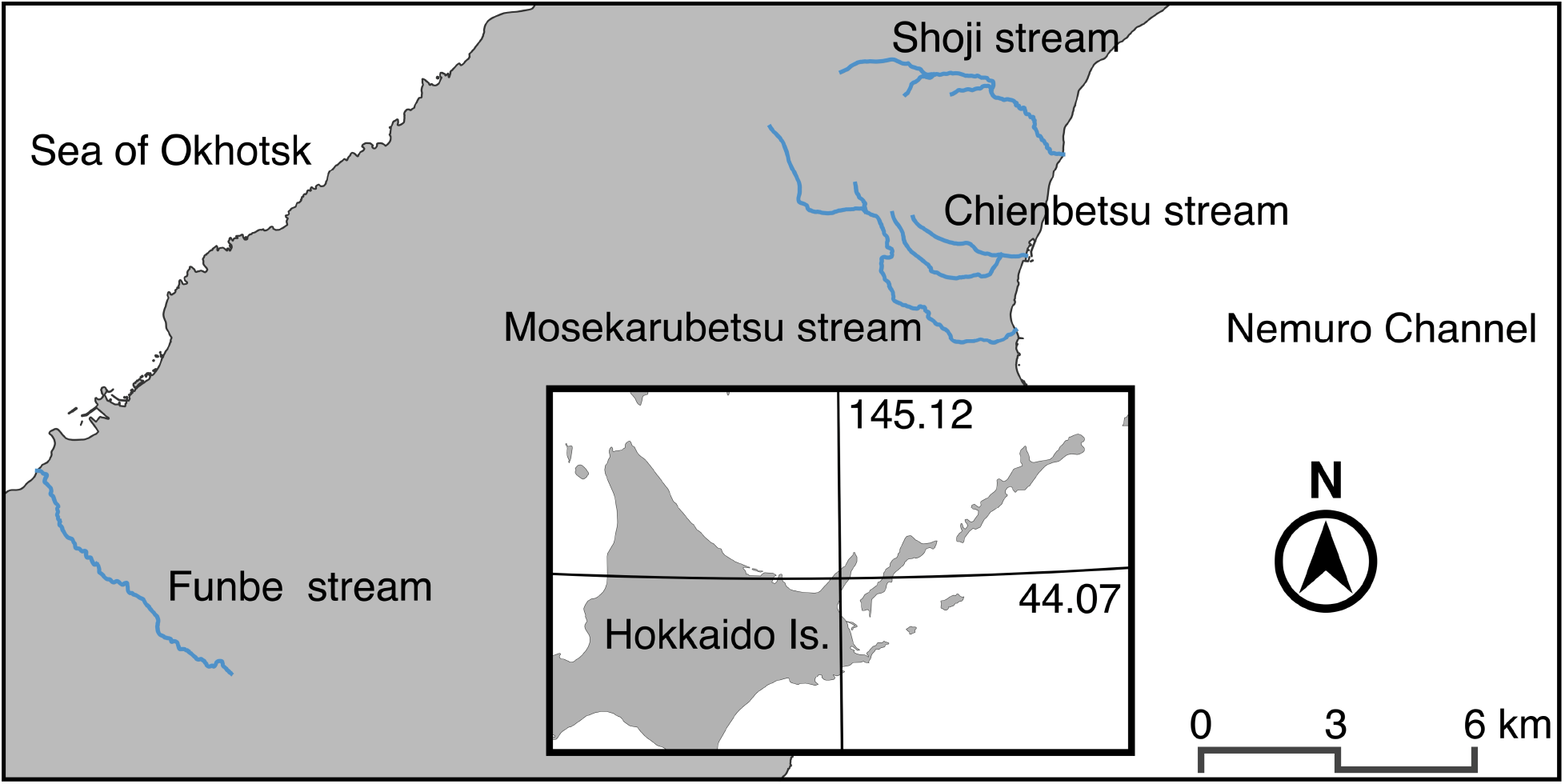
Map of the study area located on the Shiretoko Peninsula. The blue lines indicate the streams surveyed.

### 2-2. Field survey

Temporal changes in the abundance and diet of brown dippers were evaluated from mid-August to early November 2021, the spawning period of pink salmon. Field observations were conducted in one or two streams per day for a total effort of eight or nine days at every 9- to 11-day interval in each stream. In the observation protocol, one investigator (T. Yamada) walked along the study section from the lower to the upper reach, counting the number of brown dippers. To avoid recounting birds, the investigator checked where the flying individuals stopped and ignored individuals who flew ahead of the investigator, expecting territorial individuals to characteristically ‘double-back’ when pushed to the ends of their territory (Chiu et al., 2008).

The date of the count survey was the same as that of the diet survey, and the count survey was conducted before the diet survey was conducted. In the diet survey, when the investigator found an individual, they approached it at an observable distance and recorded the diet composition and age category (adult or juvenile) using binoculars (MONARCH 10 × 42; Nikon, Tokyo, Japan) (Obermeyer et al., 1999). The age category was classified by the presence or absence of juvenal plumage. The contents of the dipper’s diet were classified into four categories: aquatic insects, terrestrial insects, algae, and salmon eggs. If no observable individuals were found, no observations were made. The diet survey was conducted only once per individual at each observation cycle; the mean ± sd observation time was 4.29 ± 3.16 minutes.

Salmon spawning redds were also visually counted on the same day as the above observation procedures to obtain an index of the availability of salmon eggs. Pink salmon exhibit “probing”, a periodic and short-term migration behaviour between the sea and multiple drainages (Morita, 2021; Thedinga et al., 2000). If many individuals exhibit probing, salmon abundance cannot be a direct indicator of the number of spawners; therefore, we used the number of spawning redds as an indicator of the number of spawners. Spawning redds were visually judged as the area of disturbed gravel or bright (denuded) areas among the periphyton-covered gravel (Ortlepp & Mürle, 2003; Pedersen et al., 2009). We also measured the stream surface area of the study section only once in each section during the study period.

The abundance of brown dippers was also surveyed upstream of dams in the selected stream at the end of September 2021 during the peak spawning period of pink salmon, except in the Chienbetsu stream, where the dam has a fishway allowing migration to the upper reaches. Additional study sections were set up at approximately 400 m from the dam in each case. The distance between the end of the below dam section and the start of the above dam section in each stream was 1 to 2 m. An investigator walked from the dam to the upstream end of the study section counting the number of dippers, as was the case in the lower reach survey. This survey was conducted on the same day in the fifth cycle of the count survey mentioned above. We also measured the stream surface area of the study section only once in each stream.

### 2-3. Statistical analysis

The dependence on salmon eggs in brown dipper diets was evaluated by fitting a generalized linear mixed model (GLMM) to the individual diet data per. In the analysis, the dominance ratio of salmon eggs in the diet composition was used as a response variable and it was assumed to follow a binomial distribution. Age category and number of spawning redds were used as the candidate explanatory variables, considering stream ID and observation cycle ID as a nested random intercept. To avoid multicollinearity, variance inflation factors (VIFs) were calculated before the analysis; all variables had values less than 2.5, the threshold indicative of troubling collinearity for regressions (Johnston et al., 2018). Akaike’s information criterion (AIC) was used for model selection (Burnham & Anderson, 2002). If several plausible models had ΔAIC ≤ 2, the optimal model was selected according to the principle of parsimony (Burnham & Anderson, 2002).

A relationship between the availability of salmon eggs (represented as the number of spawning redds) and the brown dipper abundance was also estimated by fitting GLMMs to the dipper count data as a response variable assuming a Poisson distribution. The number of spawning redds and time in the day when the survey was started were used as the candidate explanatory variables, considering log-transformed stream surface area as an offset term and stream ID as a random intercept. VIF of all variables were less than 2.5. Akaike’s information criterion (AIC) was used for model selection (Burnham & Anderson, 2002). If several plausible models had ΔAIC ≤ 2, the optimal model was selected according to the principle of parsimony (Burnham & Anderson, 2002).

Since resource availability may affect organism distribution (Dingle, 2014; e.g., Dingle & Drake, 2007), it was also expected that the dipper abundance differed between the upper and lower reaches of the dam. Brown dipper abundance in the fifth observation cycle was compared between the lower reaches and the upper reaches of the dam by fitting the count data to a GLMM considering log-transformed stream surface area as an offset term and the stream ID as a random intercept. When the 95% confidence intervals (95% CI) of the estimated coefficient values did not overlap between the study sections, we considered the differences significant.

All data analyses were conducted with R v. 4.2.0 (R Core Team, 2022) using lme4 v. 1.1.30 (Bates et al., 2015) for GLMMs.

## 3. RESULTS

A total of 108 brown dipper individuals, 631 redds, 1257 pink salmon individuals, and 118 chum salmon individuals were observed during our survey. Feeding behaviour was monitored in 4 individuals (3 adults and 1 juvenile) in the pre-spawning period and in 24 individuals (15 adults and 9 juveniles) in the spawning period from the three streams, except for the Mosekarubetsu stream, where close observation could not be made.

The diet composition changed between the salmon pre-spawning and spawning periods (Figure 3a). The percentage of salmon eggs in the diet was up to 53.7% (Figure 3a) during the latter period. The brown dippers basically ingested only small food in the water and did not peck salmon carcasses. As a result of model selection based on AIC, the model including the number of spawning redds as the explanatory variable was selected (Table1), indicating that the number of spawning redds had a positive effect on the salmon egg ratio in the diet (Figure 3b; Table 2).

**Figure 3.**
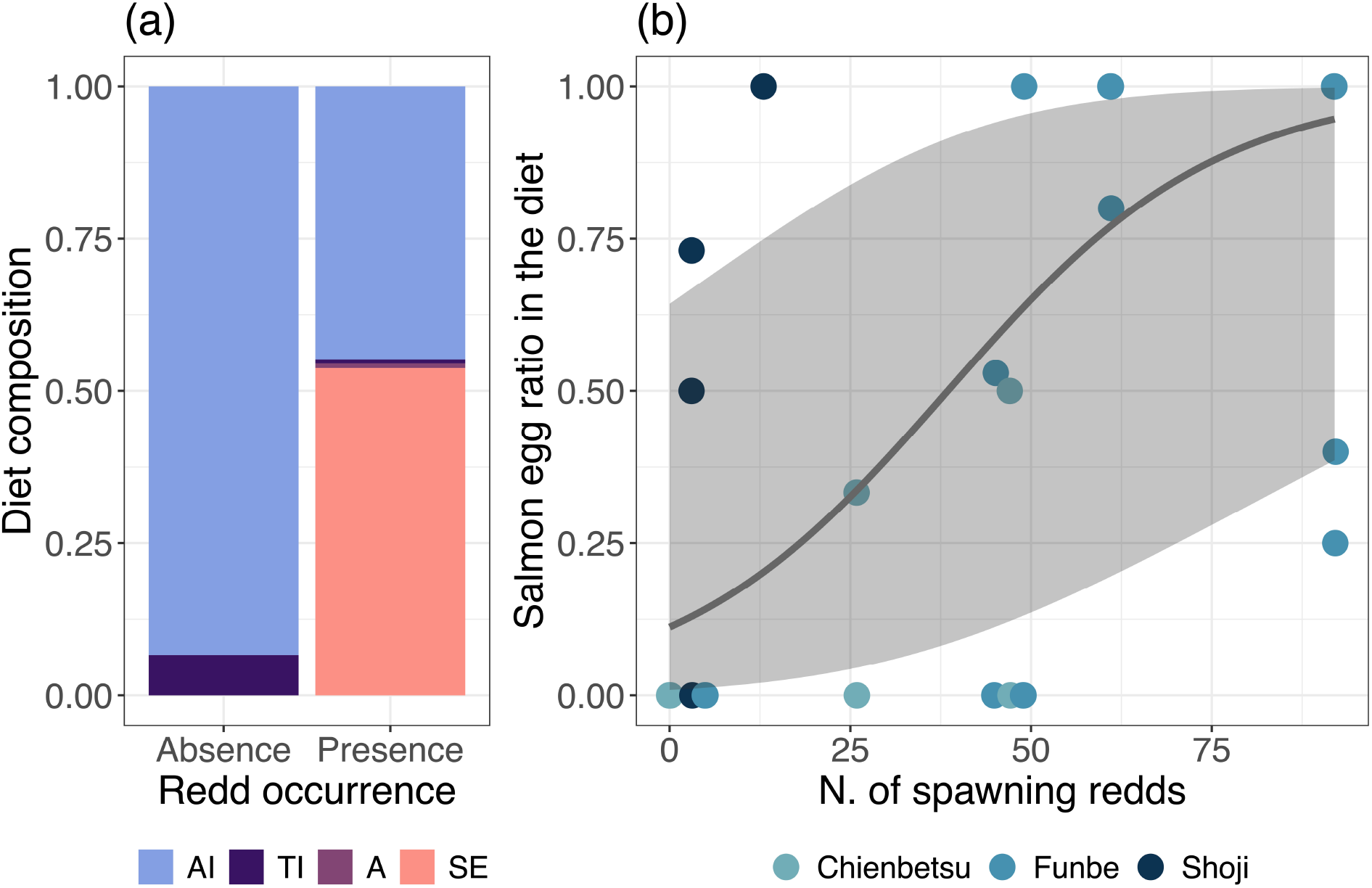
(a) Composition of brown dipper diets during the salmon pre-spawning and spawning periods. AI: aquatic invertebrate, TI: terrestrial invertebrate, A: algae, and SE: salmon egg. (b) Mean predicted marginal effects of the number of spawning redds on the salmon egg ratio in brown dipper diets. The shaded area indicates 95% CI.

**Table 1.** Results of model selection for each response variable (i.e., salmon egg ratio in brown dipper diets or brown dipper abundance). AIC corresponds to Akaike information criteria; ΔAIC is the difference between the AIC of that model and that of the model with the lowest AIC. Bold indicates the best model. Redds, number of spawning redds; Time, survey start time; Age, age category.

| Model | logLik | AIC | ΔAIC | Marginal $R^2$ | Conditional $R^2$ |
| --- | --- | --- | --- | --- | --- |
| Salmon egg ratio |  |  |  |  |  |
| **Redds** | **-45.98** | **99.97** | **0.00** | **0.319** | **0.494** |
| Redds + Age | -45.71 | 101.42 | 1.45 | 0.324 | 0.483 |
| Null | -48.91 | 103.82 | 3.85 | — | — |
| Age | -48.59 | 105.17 | 5.20 | 0.090 | 0.313 |
| Dipper abundance |  |  |  |  |  |
| **Redds** | **-63.11** | **132.23** | **0.00** | **0.396** | **0.528** |
| Redds + Time | -62.66 | 133.32 | 1.09 | 0.410 | 0.571 |
| Time | -71.60 | 149.20 | 16.97 | 0.188 | 0.311 |
| Null | -72.85 | 149.71 | 17.48 | — | — |

**Table 2.** Results of GLMMs testing effects of number of redds on the salmon egg ratio in brown dipper diets and brown dipper abundance, and the presence/absence of salmon (below/above dam) on brown dipper abundance during peak spawning period.

| Response variable | Fixed effect | Estimate | Std. Error | z value |
| --- | --- | --- | --- | --- |
| Salmon egg ratio<br>in dipper diets | Intercept | -2.07 | 1.36 | -1.53 |
|  | Number of redds | 0.05 | 0.02 | 2.70 |
| Dipper abundance | Intercept | -6.56 | 0.20 | -32.47 |
|  | Number of redds | 0.02 | 0.00 | 4.40 |
| Dipper abundance during<br>peak spawning period | Intercept | -8.32 | 0.72 | -11.52 |
|  | Section -Below dam | 2.63 | 0.75 | 3.50 |

The abundance survey showed that brown dipper abundance tended to be relatively high during the salmon spawning period in each stream (Figure 4). In the model selection process, the model including the number of spawning redds as the explanatory variable was selected (Table 1). The brown dipper abundance positively correlated with the number of spawning redds in the selected model (Figure 5; Table 2). The comparison between abundances in the upper and lower sections of the dam showed that the abundance in the lower section was clearly higher than that in the upper section (Marginal *R*^2^ = 0.792, Conditional *R*^2^ = 0.821; Figure 6; Table 2).

**Figure 4.**
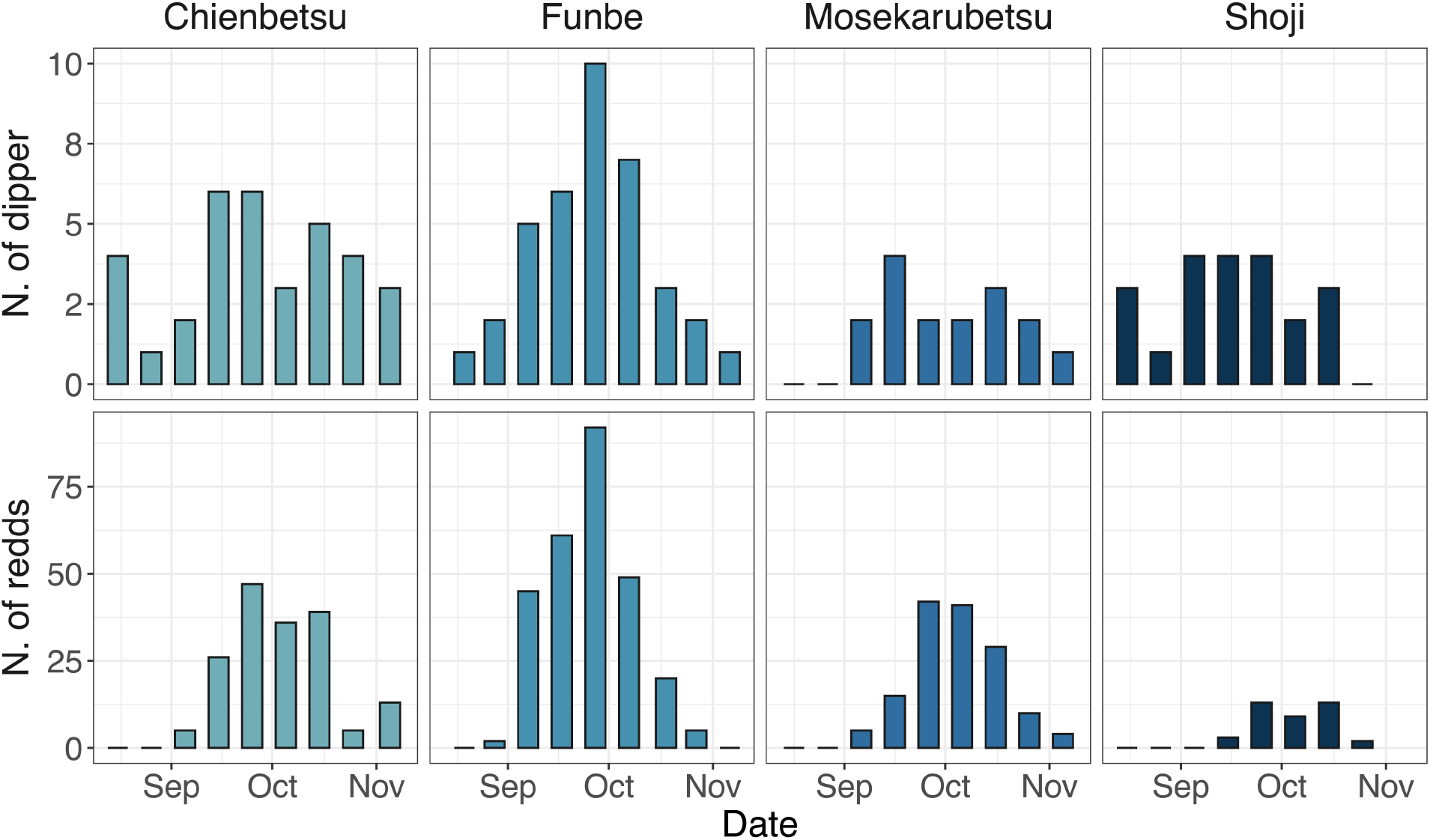
Observed brown dipper abundance and number of spawning redds in relation to surveyed date in each stream.

**Figure 5.**
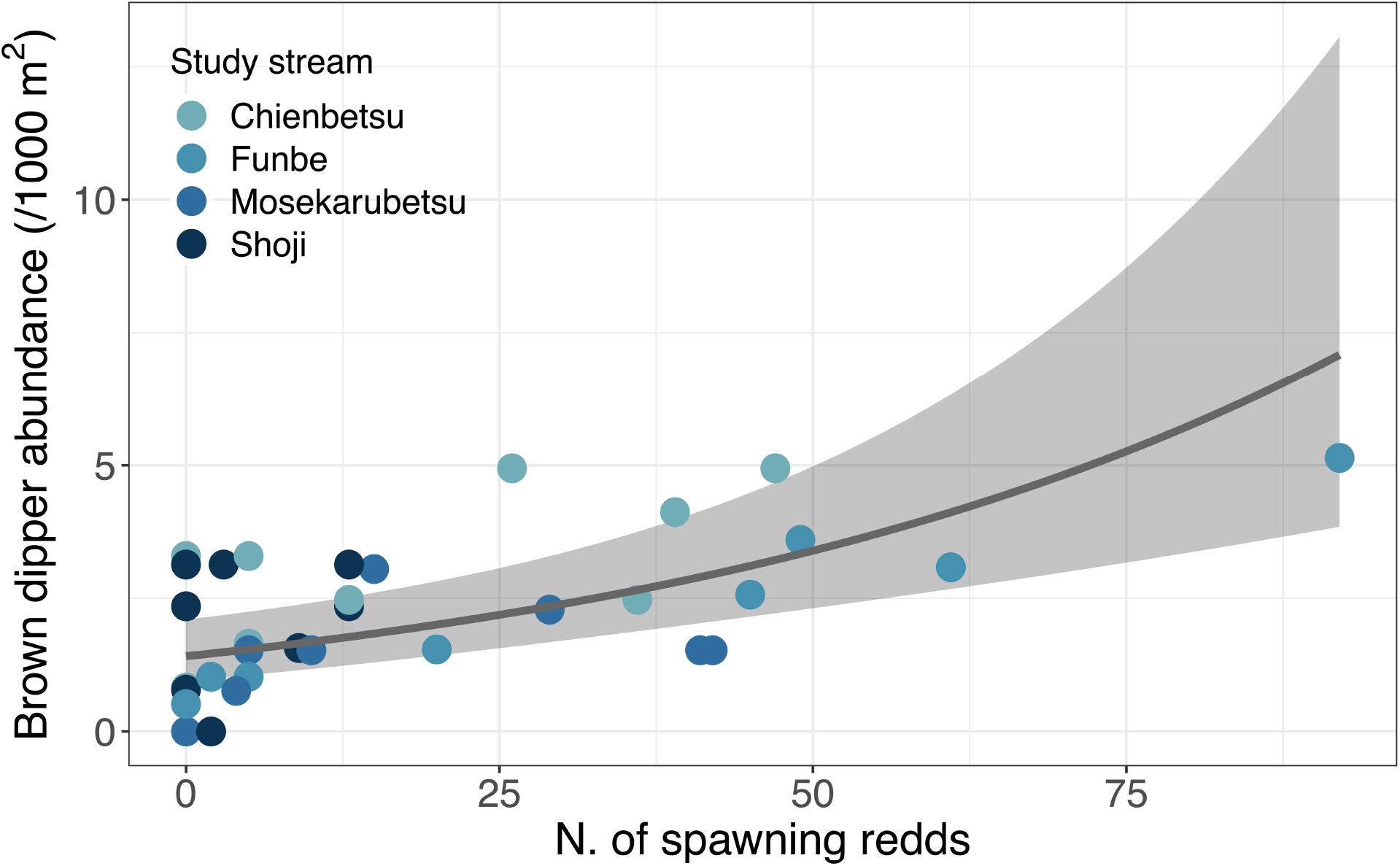
Mean predicted marginal effects of the number of spawning redds on brown dipper abundance. The 95% CI is denoted by shaded area.

**Figure 6.**
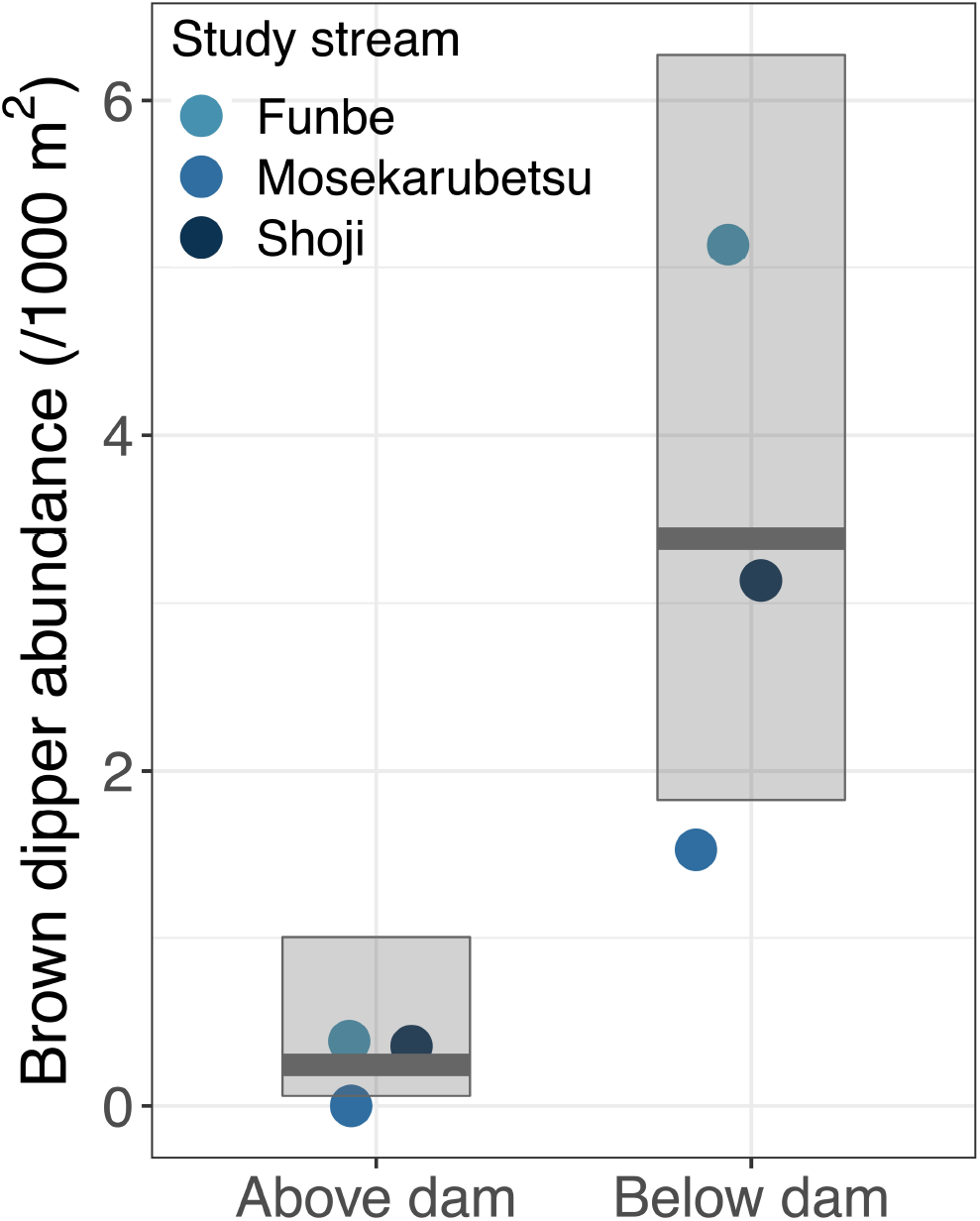
Mean predicted marginal effects of salmon occurrence on brown dipper abundance during the peak spawning season. The translucent boxes indicate 95% CIs.

## 4. DISCUSSION

This study demonstrates for the first time that salmon eggs are the dominant dietary item for the brown dipper during the salmon spawning season and that the abundance and stream distribution of terrestrial vertebrate species can be predicted by the number of spawning redds used to represent the availability of salmon eggs. In addition, brown dipper abundance during the peak spawning season differed significantly between upstream and downstream of the dam, with the downstream abundance being higher. It is therefore indicated that the distribution of brown dippers varies according to variation in the spatiotemporal availability of salmon subsidies. One of the typical theories of the spatial distribution of animal populations is the ideal free distribution, which assumes that individuals are free to move among sites (Fretwell & Lucas, 1969), and our results may be explained by this theory. The reason is that brown dippers did not use the site upstream of the dam (without salmon subsidies) even when their density was high downstream of the dam.

While salmon eggs serve as an important food source for brown dippers, the salmon spawning behaviour of digging up the riverbed leads to a reduction in the abundance of aquatic invertebrates that are prey for brown dippers (Minakawa & Gara, 2003; Moore & Schindler, 2008). Since the energy value per salmon egg is higher than that per individual aquatic invertebrate (Obermeyer et al., 2006; Whitehorne, 2010), the positive effect of egg-eating may outweigh the negative effect of reduced eating of aquatic invertebrates. In fact, juvenile weight and mortality in the American dipper in salmon spawning reaches are known to be higher and lower than those in the non-spawning reaches (Obermeyer et al., 2006). Further verification is required to clarify whether these findings may be supported in the present system.

The abundance of aquatic invertebrates varies greatly with the season (Rundio & Lindley, 2008) and declines with flooding (Chiu et al., 2008; McMullen & Lytle, 2012). Accordingly, the decrease in aquatic invertebrate abundance leads to a decline in brown dipper abundance and survival (Chiu et al., 2008, 2013). Pink salmon, chum salmon, and masu salmon running up Japanese rivers spawn during the summer, fall and winter (Iida et al., 2021; Kovach et al., 2012; Kuzishchin et al., 2009; Quinn, 2018). Since summer and fall are typhoon seasons in East Asia, salmon subsidies may compensate for the decline in aquatic invertebrates. In addition, dippers sometimes prey on salmon fry (Obermeyer et al., 2006; Ormerod, 1985; Ormerod & Tyler, 1986, 1991). Since most salmon fry mainly emerge during spring and summer (Kirillov et al., 2018; Pavlov et al., 2008; Yamada et al., 2022), salmon fry may be used as a food resource by dippers during this period. Therefore, spawning by anadromous salmonids may compensate for declines in the abundance of aquatic invertebrates during various seasons.

Salmon spawning abundance is disturbed by several human activities, such as dam construction and fisheries (Finney et al., 2000; Nakamura & Komiyama, 2010; Romakkaniemi et al., 2003). This study shows for the first time that the distribution patterns of small terrestrial predators are determined by the supply of salmon eggs, indicating that the disruption of natural spawning may have unexpected effects on the abundance and distribution of terrestrial salmon egg consumers. Future studies are needed to examine the effects of these anthropogenic restrictions of the salmon egg subsidy on the abundance and distribution of terrestrial consumers.

## ACKNOWLEDGMENTS

We thank M. Kitazawa at Hokkaido University for his valuable comments. This study was supported by JSPS KAKENHI (Grant Numbers 21H03647, 22J11475) and JST SPRING (Grant Number JPMJSP2119).

